# Ribo-attenuators: novel elements for reliable and modular riboswitch engineering

**DOI:** 10.1101/086199

**Authors:** Thomas Folliard, Barbara Mertins, Thomas P Prescott, Harrison Steel, Thomas Newport, Christopher W Jones, Travis Bayer, Judith P Armitage, Antonis Papachristodoulou, Lynn J Rothschild

**Affiliations:** Department of Biochemistry, University of Oxford, South Parks Road, Oxford OX1 3PJ, UK; Department of Engineering Science, University of Oxford, Parks Road, Oxford OX1 3PJ, UK; National Aeronautics and Space Administration Ames Research Center, Moffett Field, CA 94035, USA

## Abstract

Riboswitches are structural genetic regulatory elements that directly couple the sensing of small molecules to gene expression. They have considerable potential for applications throughout synthetic biology and bio-manufacturing as they are able to sense a wide range of small molecules and regulate gene expression in response. Despite over a decade of research they have yet to reach this considerable potential as they cannot yet be treated as modular components. This is due to several limitations including sensitivity to changes in genetic context, low tunability, and variability in performance. To overcome the associated difficulties with riboswitches, we have designed and introduced a novel genetic element called a Ribo-attenuator in Bacteria. This genetic element allows for predictable tuning, insulation from contextual changes, and a reduction in expression variation. Ribo-attenuators allow riboswitches to be treated as truly modular and tunable components, and thus increases their reliability for a wide range of applications.

## 1 Introduction

Riboswitches are structural regulatory elements generally found in the 5’ UTR of messenger RNA^1^ which allow the regulation of a downstream gene or operon in response to the binding of small molecules such as cellular metabolites or metal ions^2,3,4,5^. Relying solely on RNA for structure and appearing in all three domains of life, it is possible that they arose as an early regulatory element in a hypothesized RNA-based world^6^. Riboswitches can regulate a variety of processes including transcription, translation, and splicing in eukaryotes^7^. Riboswitches that regulate translation do so through the allosteric effects of small molecules binding to their aptamer domain. This causes a structural rearrangement which usually opens up or sequesters away a ribosome binding site (RBS) core (Figure 1A)^8,9,10^. A range of riboswitches evolved to control the expression of enzymes in natural systems, from which synthetic riboswitches have been adapted to respond to specific molecules^11,12,13,14,15,16^.

**Figure 1:**
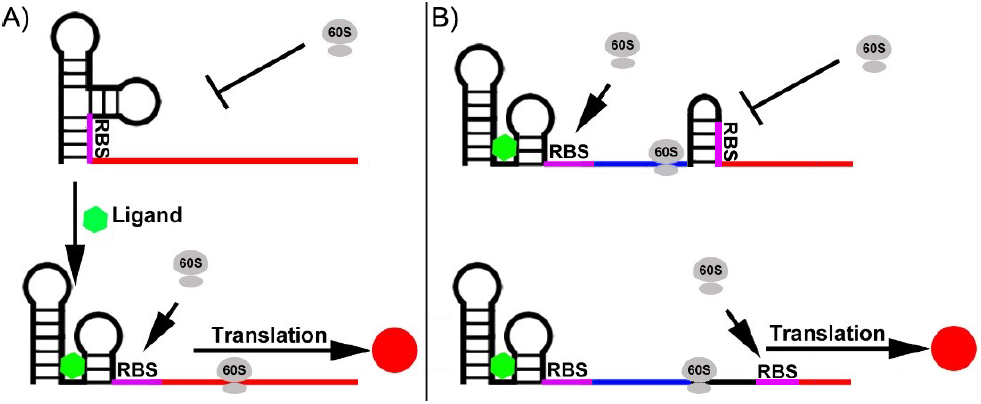
Typical activating riboswitch function: An RBS (purple) is sequestered away within a riboswitch preventing ribosome recruitment. (A) A standard Riboswitch in which binding of a ligand (green) causes a conformational change exposing the RBS, allowing translation of the gene of interest (red). (B) A Ribo-attenuator adds a second RBS, sequestered away by a local hairpin. The hairpin can be opened by a Ribosome travelling from the Riboswitch RBS, exposing the attenuator RBS and allowing translation of the gene of interest. Dynamic operation of both systems is further explained in our video animation^17^.

Ideally, riboswitches could be treated as modular “plug and play” devices to allow construction of genetic networks that respond to small molecules. However, despite significant promise and over a decade of research, limitations inherent in their function have restricted their applicability. For example, many riboswitches selectively bind a small molecule of interest using an aptamer whose specific RNA secondary structure is influenced by its own sequence, as well as the surrounding genetic context including the proximal open reading frame (ORF) under the riboswitch’s control. Thus, substituting the original ORF with a new one on a ‘start codon for start codon’ basis can nullify the desired riboswitch response to a given ligand^18^. To overcome this lack of modularity many studies have created fusions comprised of a riboswitch, the first few hundred base pairs of its working ORF, and a gene of interest^19,20,21,22^. However, this approach fails in many circumstances as it can alter the gene’s functionality, since many enzymes will not work with the inclusion of large 5’ fusions^23,24,25,26,27,28^.

Despite advances towards the *in silico* design of riboswitches^29,30,31^, adjusting their functional range in response to a given ligand remains challenging. This difficulty arises because a riboswitch’s activation or repression response is determined by both its RBS strength and secondary structure (which is itself dependent on the RBS sequence)^31^. Therefore, due to the specificity of the secondary structure surrounding the aptamer domain of riboswitch (and a poor understanding of *in vivo* RNA structures), changing the riboswitch induction range (via alteration of the RBS, generally a more predictable approach than promoter modification) without destroying the induction response is difficult. Further limitations to application of many synthetic riboswitches arise due to variation in the expression of genes under their control. This noise is thought to originate from ligand-dependent RBS accessibility bursts^8^ and limits their use with many high throughput screening techniques (such as Fluorescence Activated Cell Sorting (FACS) sorting) that rely on a distinct separation between a positive and negative population. Several solutions to these individual problems have been suggested. For example, the use of T7 polymerase could amplify a riboswitch’s output and improve its functional range^32^, however, its application is limited by its reliance on low basal expression, and does nothing to improve expression noise.

To overcome the limitations of large 5’ fusions, fixed induction ranges, and sensitivity to ORF changes of both engineered and natural riboswitches, we designed and tested Ribo-attenuators (Atts). These novel genetic elements are placed after 150 base pairs of a riboswitch’s working ORF. They consist of an RBS, over which a hairpin is engineered to silence translation independent of upstream riboswitch activity, followed by a negative one-shifted transcriptionally fused stop and start codon (TAATG) (Figure 2). The passage of ribosomes recruited by the upstream riboswitch opens up the introduced hairpin, before dissociation at the proximal in-frame stop codon (TAA) in the transcriptionally-fused junction. Additional ribosomes can then assemble at the Ribo-attenuator RBS and initiate translation at the first start codon of the introduced gene of interest (Figure 1B). Therefore, instead of directly controlling the translation of a gene as in its natural setting, the riboswitch instead controls the translational initiation rate from the downstream attenuator RBS. As such Ribo-attenuators facilitate tuning of a riboswitch’s response, the orthogonal expression of a novel gene of interest, and by improving the reliability with which the downstream RBS remains exposed, they also reduce expression variability within populations. Ribo-attenuators thus overcome many of the issues preventing wide spread use of re-engineered or synthetic riboswitches.

**Figure 2:**
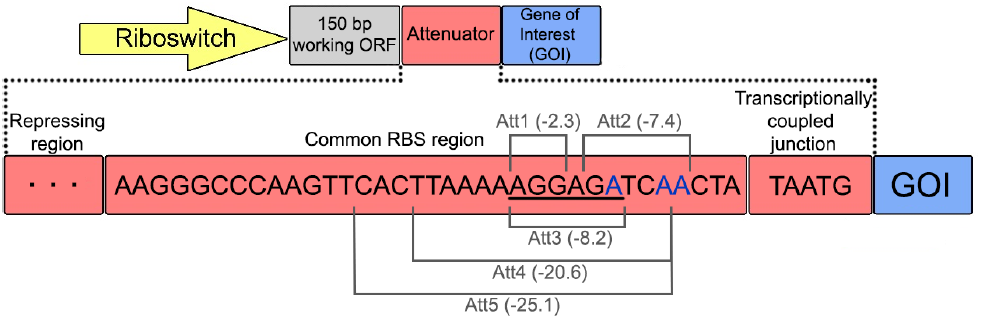
Ribo-attenuator Context and schematic: A Ribo-attenuator is inserted after a riboswitch and 150bp of its native ORF, and proceeds the desired gene of interest (GOI). We defined a Ribo-attenuator element to include a repressing region, a common RBS region (within which the RBS is underlined), and a transcriptionally coupled junction. The repressing region, which for all attenuators except Att2 is the reverse compliment of a different subsequence (indicated in grey) of the common RBS region (see Table S2 for sequences), is selected so that the attenuator region as a whole forms a hairpin with appropriate ∆G (see Figure S1 for predicted conformations). For Att2 no repressing region was included. Instead, bases highlighted in blue were altered to encourage formation of a hairpin (which involved the sequence indicated in grey) downstream of the RBS.

## 2 Materials and Methods

### 2.1 Plasmids

All riboswitch cassettes were cloned into the J64100 plasmid^33^ (ColE1 origin; 50-60 copies, Chloramphenicol resistance) under control of the tetracycline promoter. The Adda riboswitch sequence was the kind donation of Dr Neil Dixon (Department of Chemistry, University of Manchester) and the Btub riboswitch was taken from the *Escherichia coli* genome. All riboswitches were synthesised by IDT (Skokie, IL, USA) as gBlock gene fragments, assembled by Gibson assembly^34,35^ and sequence confirmed. A full list of plasmids and Addgene references is provided in Table S1. A list of Ribo-attenuator sequences is provided in Table S2.

### 2.2 Growth and Induction

All experiments were performed in *e. coli* DH5alphaZ1^36^. For each experiment three freshly transformed colonies were inoculated in LB medium (10g/L tryptone, 5g/L yeast extract, 10g/L NaCl) with chloramphenicol (25 *µ*g/mL) and grown overnight. For the functional assay and riboswitch characterisation, overnight cultures were diluted 1:100 into 1 mL LB in a deep well plate (Greiner bio-one) with Chloramphenicol (25 *µ*g/mL), Anhydrotetracycline (150 ng/mL) and stated concentrations of riboswitch inducer. For protein assays overnight cultures were diluted 1:50 into 50ml LB and the same antibiotic and inducer concentrations were used. Cultures were induced for 8 hours at 37 °C with shaking before data collection or harvesting for protein analysis. Biological triplicates were generated to assess behavioural variations between cultures.

### 2.3 Data Collection

After induction 500 *µ*L of each culture was centrifuged at 4000 RPM, washed with PBS and resuspended in 500 *µ*L of PBS. OD_600_ measurements were taken in clear well plates and GFP measurements in black well plates in a CLARIOstar platereader (BMG Labtech, Ortenberg, Germany). GFP expression was quantified by normalising by OD_600_. Flow cytometry was performed on an Attune flow cytometer (Lifetech Scientific, Basingstoke, UK) for each data point and 50,000 events were measured.

### 2.4 Fluorescence Microscopy

Fluorescence microscopy was performed on cells grown and induced as described above. Thin, flat agarose pads (1% w/v agarose in Milli-Q water) were generated on microscope slides. 2 *µ*L of cells were added to the pad immobilising them on the surface of the agarose. Cells were then imaged with a Nikon Eclipse Ti microscope (with NIS Elements software), 100× phase contrast objective (Nikon), GFP filter set (Chroma), Andor iXON CCD camera.

### 2.5 Cell disruption

Cells were harvested (3700g, 30 min) and pellets were resuspended in Lysis Buffer (PBS-Buffer with 137 mM NaCl, 2.7 mM KCl, 10 mM Na_2_HPO_4_, 1.8 mM KH_2_PO_4_, pH 7.4, with addition of Lysozyme (Sigma Aldrich), DNAse (Sigma Aldrich), and 1 protease inhibitor tablet per 50 ml solution (Complete EDTA free protease inhibitor, Roche)) relative to the wet weight of the cell pellet (1 mL buffer per 50 mg pellet). Cells were disrupted using the fast prep device (2× 6 m/s, 30s, FastPrep-24 5G, MP Biomedical) or sonication (1 min, 10s on, 4s off, 50%, Vibra-Cell VCX130, Sonics), and insoluble material was removed using centrifugation (8,000g (OmpT) or 20,000g (ColE9)), 20 min).

### 2.6 Functional assay of ColE9

Supernatants of the lysed cells were separated and diluted from 10^0^ - 10^10^ using PBS buffer. The potency was analysed by a plate killing assay^37^: An LB plate containing 0.8% Agar was warmed to 42 °C before an established culture was added in a dilution of 1:10,000. The plate was poured and left to dry for 30 min at 37 °C. 2 *µ*L of the supernatants at the given dilutions were spotted on a plate and incubated at 37 °C for 16h. The potency of ColE9 was assessed by observing the clearance around each spotted point on the plate.

### 2.7 Preparation of strep-tagged OmpT

After cell disruption, the supernatant containing membrane-bound OmpT and the pellet containing improperly folded OmpT were treated as follows. To precipitate the membranes, the supernatant was spun down (120,000g, 1.5h) and membranes were resuspended in 50 *µ*L PBS. To extract membrane bound OmpT, the resuspended membranes were diluted with 50 *µ*L PBS containing n-dodecyl-*β*-D-maltopyranoside (Anatrace) giving a final concentration of 0.635 mM (5 critical micelle concentration). The extraction was performed overnight by shaking at 4°C, and insoluble material was spun down (12,000g, 20 min) yielding soluble OmpT. The remaining pellets harbouring OmpT as inclusion bodies (IB) were washed as described previously^38^ using PBS with 1% Triton-X-100 for removal of residual *e. coli* lipids. Washed IB were dissolved using 6M Gunanidinium chloride in PBS and incubated for 10 min at 95°C. Insuloble material was removed (12,000g, 5 min) yielding the IB fraction. Whole cells (WC), extracted membrane bound OmpT (Mem) and solubilized misfolded OmpT (IB) were resolved on 4-12 gels [Q][what’s a 4-12 gel?], using the Laemmli buffer compositions^39^. Immunoblotting was performed as described previously^40^ using anti-strep antibodies (Thermo Fisher Scientific) and Luminata Western HRP Substrate (Merck Millipore).

### 2.8 Model Methods

The cumulative distribution of the random variable T denoting the time elapsed between an arbitrary time-point T and the next production of a GFP molecule was calculated according to the model described in the Supplementary Information. This model treats the transition between closed (OFF) and open (ON) states of both the riboswitch and attenuator as stochastic random processes, with the rate of attenuator opening taking an increased value when the riboswitch is ON (Figure 6A). This dependency captures the system’s intended mechanism, whereby a ribosome initiated at the riboswitch RBS opens the attenuator, revealing the second RBS. The primary assumptions made are therefore the model’s parameterised rate values, for which we used the following:

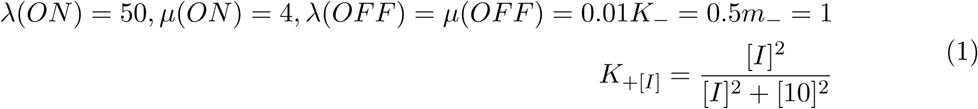

For 0 ≤ [*I*] ≤ 50 (where [*I*] = 50 corresponds to 100% induction). The inverse of T is the random variable we call “Expression Rate”. For the calculation of the CDFs, see Supplementary Information and the corresponding MATLAB code.

## 3 Results

### 3.1 Change in open reading frame demonstrates sensitivity of 5’ regulatory elements to secondary structure

Two previously reported riboswitches were investigated for sensitivity to changes in ORF. The Adda riboswitch, characterised from *Vibrio vulnificus* ^15,41^, is an activating purine riboswitch^42,43,44^ reported to selectively bind 2-aminopurine. Fusing sfGFP to the first 150 base pairs of the previously reported working ORF^15^ yielded an induction response (to 2-aminopurine concentrations between 0 and 250 *µ*M) very similar to previously reported studies. However, directly replacing the reported ORF with sfGFP (start codon for start codon) gave no induction response. These data were supported by single cell analysis, which demonstrated a response to the inducer for the fusion construct, but no such output when the ORF was completely substituted (Figure 3A).

**Figure 3:**
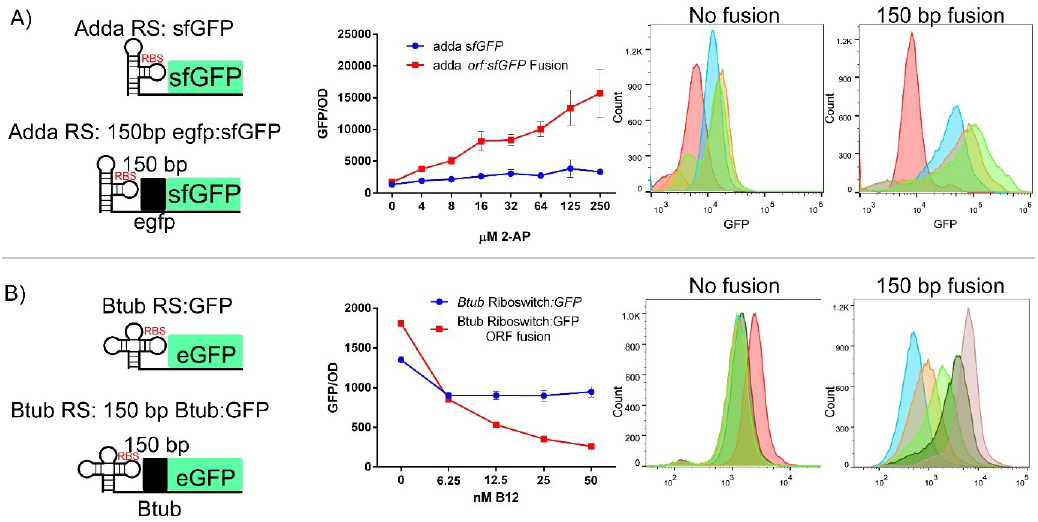
Adda and Btub riboswitches exhibit contextual sensitivity: (A) The Adenine riboswitch had sfGFP Introduced directly after the riboswitch replacing its reported ORF (previously eGFP), and also fused to the first 150bp of the previous ORF. Both constructs were analysed for population and single cell response to 2-aminopurine. Error bars indicate standard deviation around the mean for biological triplicates. Single cell colours; Red (0 *µ*M) Blue (8*µ*M) Orange (32*µ*M) Green (250*µ*M). (B) The Btub riboswitch had eGFP introduced directly after the riboswitch replacing its reported ORF (previously Btub), and also fused to the first 150bp of the previous ORF. Both constructs were analysed for population and single cell response to adenosylcobalamin. Single cell colours; Red (0 nM) Dark Green (6.25 nM) Light Green(12.5 nM) Orange (25 nM) Blue (50 nm).

The Btub riboswitch is a repressive riboswitch notable for its response to the high-value small metabolite adenosylcobalamin, an active form of vitamin B_12_ ^19^. As with the Adda riboswitch, replacing the working ORF with eGFP (start codon for start codon) resulted in a loss of induction response (between 0 and 50 nM of adenosylcobalamin) for both colony and singlecell measurements. However, fusion of eGFP to 150 base pairs of the working ORF yielded the desired repressive induction response (Figure 3B).

### 3.2 Design of Ribo-attenuators

Ribo-attenuators were designed within the Adda riboswitch system, placed between 150bp of the system’s previous working ORF (eGFP) and the new gene of interest (sfGFP), as illustrated in Figure 2. First, a common RBS region was designed to include the SD1 RBS from the BIOFAB parts library^45^, which was placed upstream of a transcriptionally coupled junction (TAATG) that included a stop codon in-frame with any ribosome originating from the riboswitch, and a negative one base shifted start codon. Hairpins with ever lower ∆G were introduced over the RBS on the 5’ end by altering the attenuator’s repressing region (Figure S1). For one attenuator (Att2) we took a different approach, engineering a hairpin on the 3’ side by altering bases within the common RBS region. Screening of hairpins with stems of 3-24 bp in length yielded a step-like expression profile with intervals of 3 bp, from which we selected the least variable Ribo-attenuators at each plateau. Further Ribo-attenuators could easily be designed rationally around a new RBS by following our process, or by generating random sequences and screening for a Ribo-attenuator of the required strength. The selected Ribo-attenuators (without upstream riboswitch) were expressed under the control of the tetracycline promoter, demonstrating that with the exception of the smallest hairpin (Att1), the hairpin structures caused a significant drop in the RBS’s translational efficiency (Figure S2). Thus, when a riboswitch is placed upstream of the Ribo-attenuator translation from the attenuator’s RBS independent of riboswitch activity will be minimal.

### 3.3 Ribo-attenuators can shift Adda riboswitch induction response

Five Ribo-attenuators were introduced downstream of the adenine riboswitch and 150 bp of its working ORF (eGFP). The Ribo-attenuators were able to shift the system’s induction profile in a manner that correlated with the strength of the attenuator; a stronger hairpin leads to a narrower and lower induction response (Figure 4A). The direct fusion of sfGFP to the working ORF yielded very clear inclusion bodies manifest as distinct spots present at one pole of the cell (Figure 4B), which are formed by misfolded insoluble proteins. By comparison, the inclusion of a Ribo-attenuator between the riboswitch and sfGFP yielded soluble protein (Att1 shown) since the translated reporter gene is expressed orthogonally from the upstream 150 base pairs of the working ORF. The absence of the fusion domain in the Ribo-attenuated case was confirmed via immunoblotting, which demonstrated a size difference between the protein products in systems with and without the attenuator (Figure S3).

**Figure 4:**
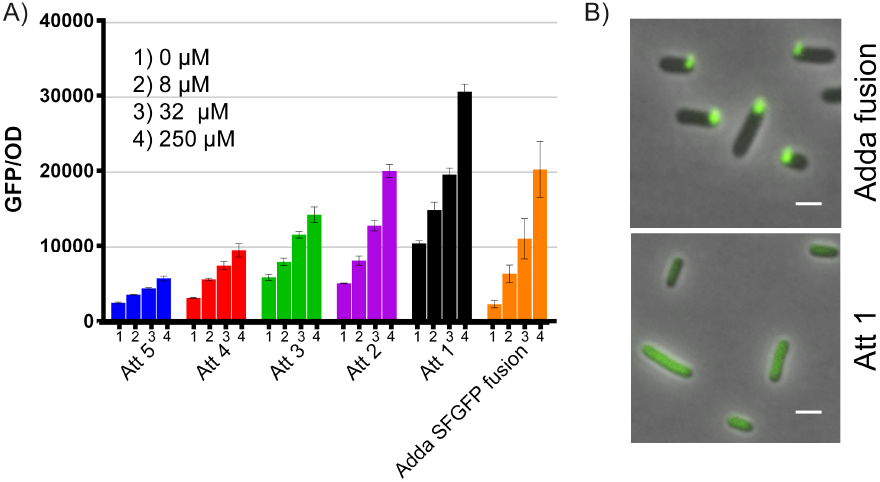
Ribo-attenuators allow Adda riboswitch rational re-engineering. (A) Response of each Ribo-attenuator to 0, 8, 32 and 250 *µ*M 2-aminopurine as compared to the fusion construct (orange). Error bars indicate standard deviation around the mean for biological triplicates. Full single-cell data is presented in Figure S5. (B) Fluorescence microscopy of the Adda fusion showing clear inclusion bodies and the Att1 Ribo-attenuator showing soluble dispersed GFP. White bars represent 1*µ*m.

To demonstrate the universal applicability of Ribo-attenuators, the Adda riboswitch was used in isolation, as a 150 bp fusion with its working ORF (eGFP), and with Att2, to express and analyse the targeting of OmpT, an outer membrane protein. For all constructs basal expression of OmpT was observed (Figure S4). The riboswitch in isolation failed to produce membrane bound OmpT, whilst the fusion system yielded mainly insoluable inclusion bodies. However, the Ribo-attenuated system yielded correctly targeted OmpT in the cell membrane, demonstrating the Ribo-attenuator’s ability to express working protein.

### 3.4 Ribo-attenuators can shift and amplify Btub riboswitch induction response

The same five Ribo-attenuators were introduced downstream of the Btub riboswitch and 150 bp of its working ORF (Btub). The Ribo-attenuators were able to shift and substantially amplify the system’s induction profile, with (as for the Adda riboswitch) increased hairpin strength corresponding with reduced expression (Figure 5A). Introduction of the attenuators resulted in a notable difference between the activated and unactivated systems when observed by fluorescence microscopy (Figure 5B). The use of the same five Ribo-attenuators demonstrates the modularity of our system: Despite these attenuators being designed within a different genetic context (the Adda riboswitch system), when transplanted into the Btub system their functionality is maintained.

**Figure 5:**
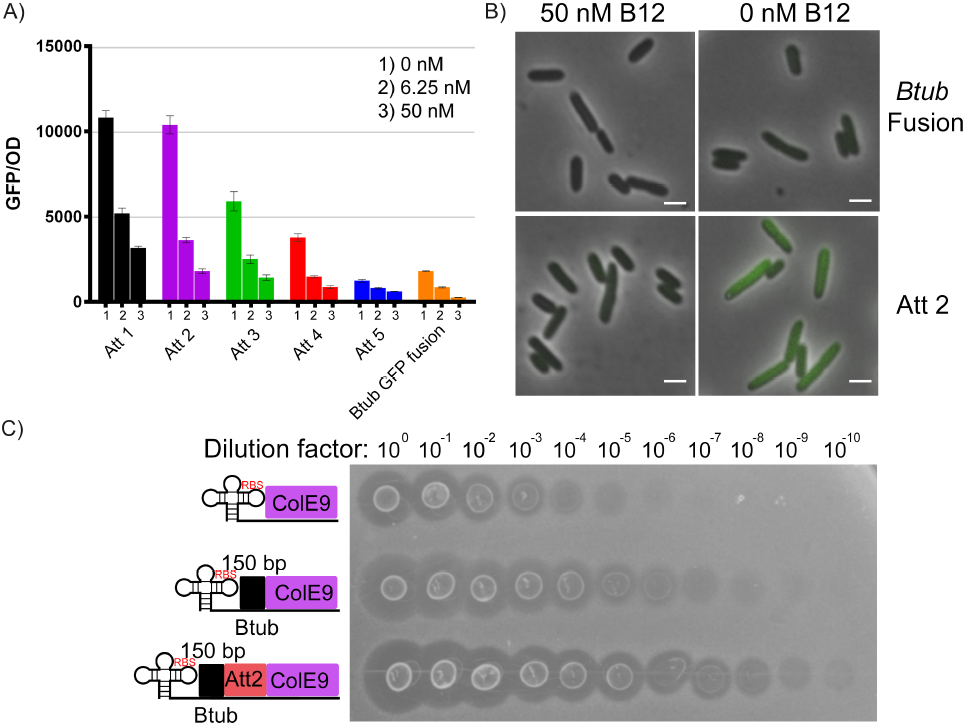
Ribo-attenuators allow Btub riboswitch rational re-engineering. (A) Response of each Ribo-attenuator to 0, 6.25, and 50 *µ*M 2-aminopurine as compared to the fusion construct (orange). Error bars indicate standard deviation around the mean for biological triplicates. Full single cell data is shown in Figure S5. (B) Fluorescence microscopy of the uninduced and induced Btub fusion and Att2 ribo-attenuated Btub riboswitch, with the latter demonstrating a greater induction difference. White bars represent 1*µ*m. (C) Plate-killing assay for production of bacteriocin ColE9 from the Btub riboswitch with and without fusion and attenuator domains. Clearances in the *e. coli* soft agar plate correlate with ColE9 production level, demonstrating a substantially increased protein yield in the Ribo-attenuator system.

**Figure 6:**
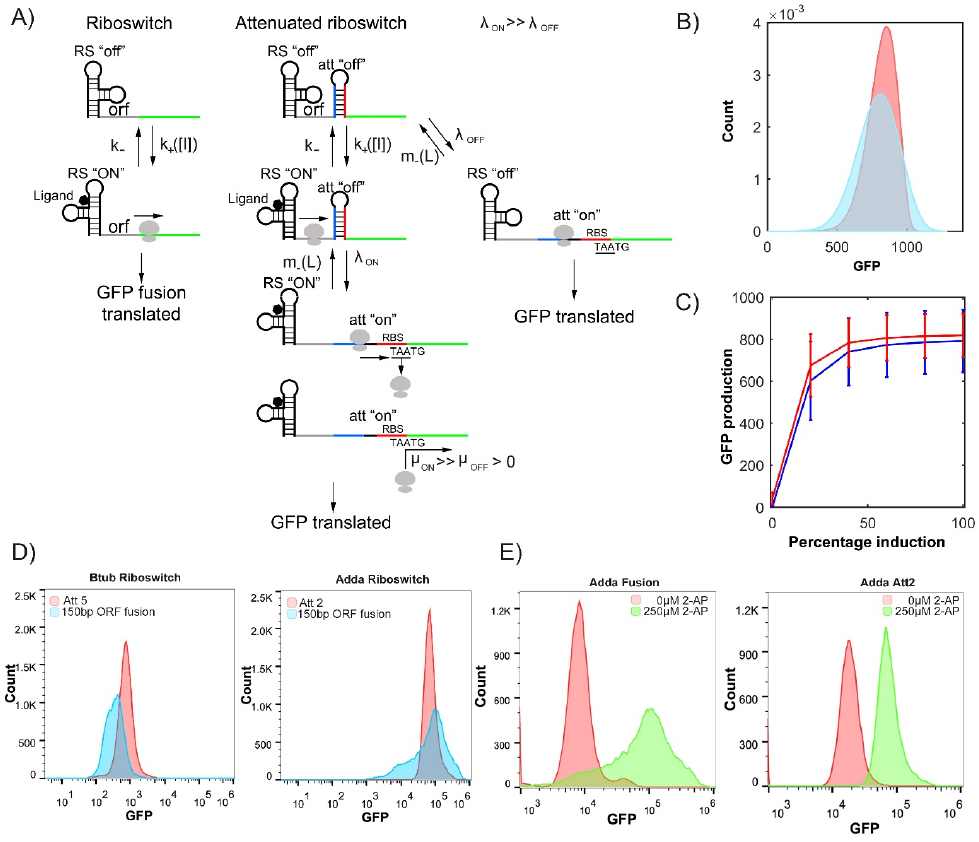
Ribo-attenuators reduce riboswitch expression noise. (A) Random walk dynamics of the modelled system. The riboswitch of the one-component system randomly switches between OFF and ON states at rates k_+_[I] and k_−_, and GFP is produced at rate *λ*_*OFF*_ or *λ*_*ON*_ depending on the state of the riboswitch. In the two-component (Ribo-attenuator) system, the riboswitch has similar dynamics; the rates *λ*_*OFF*_ and *λ*_*ON*_ are now the rates at which the downstream attenuator is switched ON. The downstream attenuator can spontaneously switch off at rate m_−_(L), which is an increasing function of the length L of the attenuator’s hairpin stem. GFP is produced at rates *µ*_*OFF*_ and *µ*_*ON*_ depending on the state of the downstream attenuator. (B) Probability distribution for the random variable defined as the number of GFP molecules expressed by the stochastic model in (A) over a unit time interval starting from an arbitrary (long-term) time-point at maximal (100%) induction for the one-component riboswitch system (blue) and the attenuated system (red). (C) The expression distribution mean plus/minus one standard deviation (error bars) for varying induction concentrations defined as a percentage of maximal induction (the data points at 100% induction therefore corresponds to the data in (B)). Comparison of the one-component riboswitch (blue) and Ribo-attenuator system (red) demonstrates substantially reduced variability across the induction range. (D) Measured expression distributions at maximal induction for the Btub riboswitch with a single-component fusion system (blue) and with Att5 (red), and for the Adda riboswitch with a single-component fusion system (blue) and Att2 (red). (E) Single cell expression distributions at the lower and upper limits of the Adda riboswitch system’s induction with and without inclusion of the Att2 Ribo-attenuator. Equivalent data for the Btub system is presented in Figure S6.

To demonstrate further application of Ribo-attenuators, the Btub riboswitch was used in isolation, as a 150 bp fusion with its working ORF (Btub), and with Att2, to express ColE9, a bacteriocin that invades (via OmpF and BtuB) and attacks the genome of *e. coli* cells^46^, preventing their growth. The amount of active ColE9 in a sample can be directly correlated with the clearance of *e .coli* grown on a soft Agar plate when ColE9 is spotted on to it^37^. As in Figure 3B, it was found that inserting ColE9 directly after the riboswitch resulted in minimal expression (Figure 5C). Expression was increased by a factor of ~ 1000 in the fusion system, however, with the inclusion of Att2 a substantially higher yield of active protein was achieved: Clearances were observable up to dilution by a factor of 10^10^, demonstrating the attenuator’s ability to significantly enhance protein production from this riboswitch.

### 3.5 Ribo-attenuators reduce uncertainty of riboswitch function

The translation of GFP from mRNA in the modelled one-component riboswitch system at any given time depends on the state of the riboswitch, either *λ*_*ON*_ or *λ*_*OFF*_ (Figure 6A). Over time the riboswitch stochastically switches between ON and OFF, with inducer concentration biasing the stochastic switching towards one or the other state. Assuming that the two translation rates are at different orders of magnitude, the waiting time from any given time until the production of a GFP molecule is thus highly uncertain. In the two-component case (with the addition of a ribo-attenuator), the accessibility of the two RBSs can be in one of four states (OFF, OFF), (OFF, ON), (ON, OFF), and (ON, ON) (Figure 6A). The rates *λ*_*ON*_ and *λ*_*OFF*_ of ribosome transit now determine the rates at which the second switch turns on, given the state of the first. The scale separation of those rates means that, to a first order approximation, the first switch ON implies the second switch will turn ON, and the first switch OFF implies the second switch is likely to turn OFF. However, the GFP translation rates *µ*_*ON*_ and *µ*_*OFF*_ resulting from the state of the second switch can now be set independently of the translation parameters of the upstream riboswitch via adjustment of the Ribo-attenuator’s RBS. Expression level is also dependent upon the rate at which the Ribo-attenuator switches back to the OFF position (m_−_(L)), which is itself a function of the length (L) and hence strength of the attenuator region’s secondary structure.

From this stochastic model probability distributions for total GFP expression were generated, demonstrating a substantial reduction in gene expression variability (at maximal induction) for the Ribo-attenuator system when compared to the un-attenuated riboswitch (Figure 6B), a trend which is maintained over a range of induction levels (Figure 6C). The same behaviour was observed experimentally for both the Adda and Btub riboswitches (Figure 6D), which demonstrate that inclusion of a Ribo-attenuator substantially reduces the width of the expression distribution (though this comparison relies on the assumption that unpredictability in GFP expression is a direct consequence of uncertainty in GFP translation initiation rates [is this caveat necessary to include?]). Figure 6E displays the single cell distinction between uninduced and fully induced populations for the Adda riboswitch with and without Ribo-attenuator, demonstrating that the reduced variation in the attenuated system makes overlapping populations substantially more distinct. A similar trend was observed for the Btub riboswitch (Figure S6).

## 4 Discussion

We designed a novel and widely applicable tool that allows riboswitches to be used as modular, tunable components with a highly predictable outcome, enhancing applicability by overcoming previously identified drawbacks. We chose two well known, representative riboswitches; the Adda riboswitch, an activating class of riboswitches that respond to a cellular metabolite (Adenine) and the Btub riboswitch, a repressing riboswitch which responds to adenosylcobalamin. These have the potential to facilitate the biological production of high value micronutrients such as vitamins, many of which are still produced via expensive and inefficient chemical synthesis. Recently described methods applying riboswitches to high value vitamins could also be used with the Ribo-attenuators described herein to elicit a high throughput approach to pathway elucidation^47^.

The first advantage of the developed Ribo-attenuator system is its enabling of riboswitch controlled expression of a new gene of interest without the inclusion of a 5’ fusion. Typically such fusions have been employed^19,20,21,22^ because direct replacement of a riboswitch’s ORF can degrade its functionality^18^ (as demonstrated in Figure 3). However, re-engineering a riboswitch in this fashion gives rise to a protein with a large fusion, which can severely impact function. For example, this was demonstrated in in Figure 3B, in which the fusion approach led to formation of inclusion bodies. As a second example, expression of the outer membrane protein OmpT demonstrated that fusion to the first 150 bp of Btub resulted in formation of inclusion bodies, whereas the attenuator-produced protein was correctly targeted to the membrane. We hypothesise that inclusion bodies are present because the riboswitch’s working ORF contains a beta strand, which when expressed as a fusion causes mis-folding due to the resultant protein having an uneven number of beta strands. Additionally, in the case of OmpT the fusion domain’s presence may disrupt the signal sequence required for membrane transport. Ribo-attenuators overcome this challenge by translating a gene of interest from a second downstream RBS, avoiding inclusion of any 5’ fusion and thereby improving functionality (Figure 3B).

A second benefit provided by inclusion of a Ribo-attenuator is the ability to modularly tune the induction response of a riboswitch. This was demonstrated by Ribo-attenuators shifting and re-scaling the output response of the Adda system in which they were designed (Figure 4A), in a manner correlating with *a priori* predictions of their relative strength. This functionality was maintained when the same Ribo-attenuators were employed in a different genetic context in the Btub system (Figure 5A), demonstrating their modularity. In both contexts an increased (compared to the fusion system) expression level was observed when the riboswitch was in its repressing induction state. In addition to spontaneous accessibility of the Ribo-attenuator’s RBS, this may arise since any ribosome originating from the riboswitch RBS would in the fusion case produce a single copy of GFP, whilst in the Ribo-attenuator system it would open the attenuator’s hairpin for a time period during which multiple translation initiation may events occur. Particularly in the Btub system Ribo-attenuators were able to substantially increase the maximal induction output (for both GFP and ColE9). This is an important capability^32^ that would be difficult to achieve via re-engineering of the riboswitch’s RBS (which could influence its ligand-binding ability), or via increasing translational promoter strength (our system already employed the strong tetracycline promoter at full induction). Though attempts have been made to overcome these problems, such as boosting riboswitch efficiency through RNA amplification^32^ or de-novo design of synthetic riboswitches^31^, as-yet they do not provide a reliable, modular, and predictable approach to tuning of riboswitch induction. This is achieved by our Ribo-attenuator system, for which new variants can be developed with different RBS strengths and easily predicted secondary structures, whilst preserving riboswitch ligand response.

A third advantage provided by Ribo-attenuators is their ability to reduce variability of a riboswitch’s output. This limitation had restricted the application of riboswitches in a number of ways: For example, unpredictable behaviour may produce downstream variance in processes under a riboswitch’s control, whilst also making separations of populations by FACS challenging, limiting the ability to screen libraries for novel compounds. Our modelled Ribo-attenuator system provided a theoretical underpinning for its noise-reduction capability, demonstrating that inclusion of a Ribo-attenuator can cause a substantial reduction in population expression variability, as well as a slight increase in mean expression (Figure 6B). This was observed experimentally (Figure 6D), demonstrating that the reduction in variability can provide more distinct output distributions (Figure 6E).

In conclusion, by fine tuning sensitivity, and reducing uncertainty in expression thereby separating cell populations, Ribo-attenuators facilitate the use of riboswitches in applications ranging from FACS screening of component libraries to rapid elucidation of novel biosynthesis pathways. Initially we examined the sensitivity of two well-known riboswitches to the introduction of a new open reading frame: Introducing a novel reporter as part of a fusion yielded functional riboswitches, but inclusion bodies that significantly limit application were observed. We also found a high degree of variation in the Adda riboswitch, which made identification of specific concentrations of ligand difficult. Using Ribo-attenuators we allowed both riboswitches to maintain the original open reading frame and as a result their ligand response, whist expressing the introduced gene of interest orthogonally over a predictable dynamic range. Using these Ribo-attenuators it should be possible to overcome limitations to riboswitch identified in previous studies. For example, the Glycine riboswitch is a similar riboswitch from *Bacillus subtilis* identified as having a narrow induction range^48^, which limits its use as a tool for maintaining vectors in the absence of an antibiotic, or as a cheap Glycine-based induction system for bioreactors. As shown in Figure 5 for the Btub riboswitch, Ribo-attenuators could amplify and tune this system’s static response, aiding its industrial application. In summary, Ribo-attenuators represent a breakthrough step towards allowing riboswitches to be treated as truly modular and tunable devices.

## Acknowledgments

Drew Endy, George Wadhams, Chris Grant, Ciarán L Kelly, Karl Brune, Axel Nystrom. Members of the Bayer and Papachristodoulou groups.

### Funding

Support for this work came from the EPSRC project EP/M002454/1. and the BBSRC project BB/L002507/1

### Author Contributions

TF, TB, JPA, TN and LJR designed the research, TF, BM and CWJ performed experiments, TPP, HS and AP did modelling and computational work. All authors analysed data and wrote the paper.

### Competing financial interests

The authors declare no competing financial interests

